# Mixture-of-Experts Approach for Enhanced Drug-Target Interaction Prediction and Confidence Assessment

**DOI:** 10.1101/2024.08.06.606753

**Authors:** Yijingxiu Lu, Sangseon Lee, Soosung Kang, Sun Kim

## Abstract

In recent years, numerous deep learning models have been developed for drug-target interaction (DTI) prediction. These DTI models specialize in handling data with distinct distributions and features, often yielding inconsistent predictions when applied to unseen data points. This inconsistency poses a challenge for researchers aiming to utilize these models in downstream drug development tasks. Particularly in screening potential active compounds, providing a ranked list of candidates that likely interact with the target protein can guide scientists in prioritizing their experimental efforts. However, achieving this is difficult as each current DTI model can provide a different list based on its learned feature space. To address these issues, we propose EnsDTI, a Mixture-of-Experts architecture designed to enhance the performance of existing DTI models for more reliable drug-target interaction predictions. We integrate an inductive conformal predictor to provide confidence scores for each prediction, enabling EnsDTI to offer a reliable list of candidates for a specific target. Empirical evaluations on four benchmark datasets demonstrate that EnsDTI not only improves DTI prediction performance with an average accuracy improvement of 2.7% compared to the best performing baseline, but also offers a reliable ranked list of candidate drugs with the highest confidence, showcasing its potential for ranking potential active compounds in future applications.

**CCS CONCEPTS:** • **Applied computing** → **Bioinformatics**; • **Computing methodologies** → **Artificial intelligence**.

## 1 INTRODUCTION

In recent years, AI-driven drug discovery has surged due to the growth of deep learning algorithms and biomedical data. Unlike traditional *in vitro* processes, which are time-consuming and costly, computer-aided drug discovery (CADD) has become crucial for accelerating drug discovery [1]. A key focus within CADD is accurately identifying complex drug-target interactions (DTI), a pivotal step in drug development [2]. Advanced models like ChemBERTa [3] for molecule prediction, ProtBERT [4], ESMFold [5] for protein structure, and AlphaFold 3 [6] have ushered in a new era of AI-driven drug discovery, capable of high-precision predictions. However, the limited daily access, high inference costs, or extensive inference times still pose significant challenges for researchers aiming to screen vast libraries of drug candidates [7–9]. Thus, smaller deep learning models are preferred for drug candidate screening [10, 11]. These models, trained for DTI tasks, leverage diverse pharmacological, genetic, and sequential features to learn latent embedding spaces with state-of-the-art techniques, demonstrating expert-level performance across numerous datasets [12**–** However, Despite the significant advancements achieved by deep learning-based DTI models, they still face critical challenges related to limited confidence estimation and a lack of consensus [17]. During experiments with some existing deep learning-based models (Figure 1), we observed the following phenomena: (1) These DTI expert models exhibit a lack of consensus on the same test data, producing divergent outcomes for identical test samples. (2) The performance of the same expert varies across different datasets. These inconsistencies make it challenging for researchers to discern which model predictions are the most reliable for downstream drug development processes. Furthermore, most existing models do not offer a measure of the reliability of their predictions, leaving researchers uncertain about the extent to which they can trust and apply these predictions in practical drug development for potential active compound screening. Providing a reliable ranking list of compound candidates for specific target proteins can help scientists prioritize their experimental efforts.

**Figure 1:**
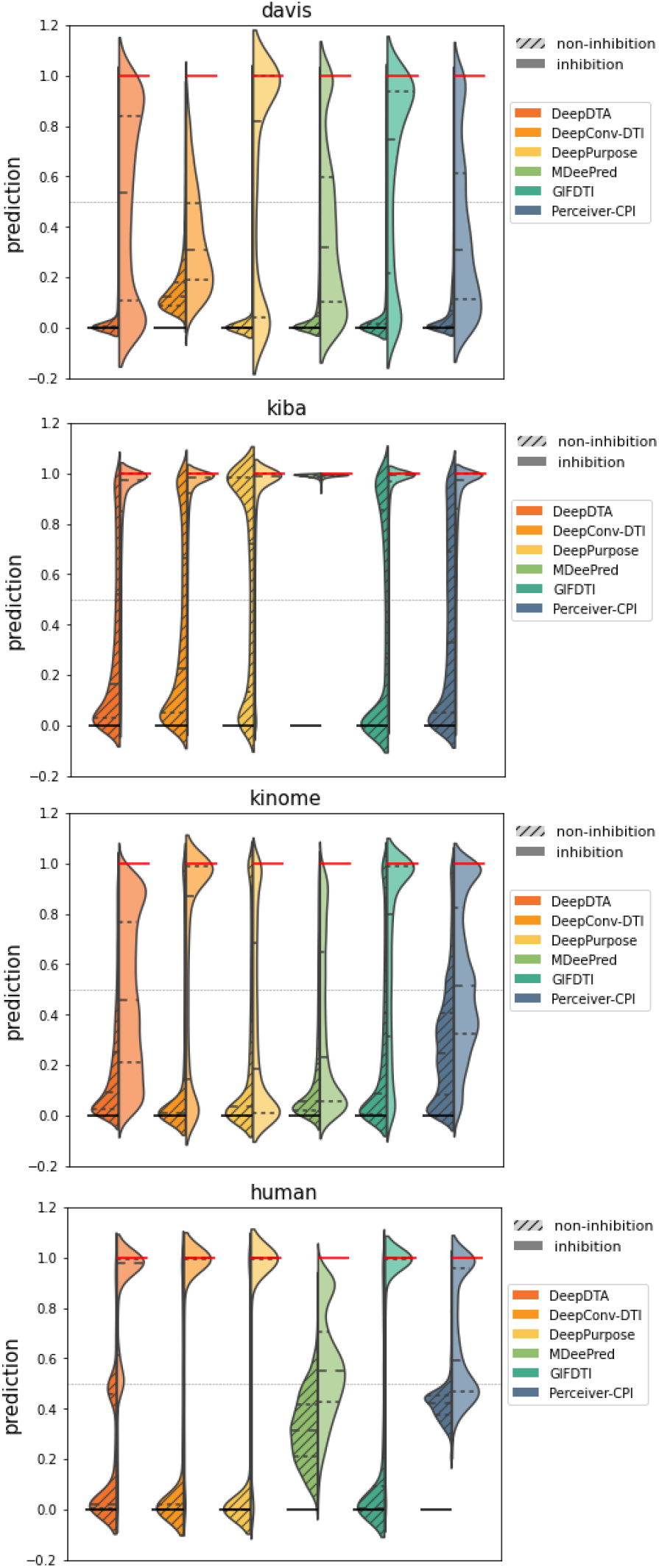
Distribution of predictions over different databases made by current DTI experts. We draw the distribution of prediction of each DTI expert model under different database to see whether there is any superior model over databases. Left and right portion of violin chart represent the prediction distribution of data with the true label of positive and negative respectively.

To address these challenges, we propose EnsDTI, a mixture-of-experts (MoE) approach designed to enhance the prediction accuracy and reliability of existing expert models across different data distributions. EnsDTI follows a four-stage training process. Initially, each DTI expert model is trained to achieve its maximum performance. Subsequently, we freeze all the expert models and generate their prediction probabilities. After, we train a gating network based on their predictions to integrate the knowledge from these experts. In the final stage, we apply an independent inductive conformal predictor module to assign confidence values to the predictions of EnsDTI, thereby providing a reliable ranking list of candidate drugs. Empirical experiments on four popular benchmark datasets demonstrate that EnsDTI effectively improves drug-target interaction predictions compared to current DTI expert models. Case studies further reveal that our model exhibits superior consistency and accuracy in predicting probable candidate lists for different kinase proteins with similar functions.

## 2 RELATED WORKS

Accurately identifying complex interactions between drugs and targets is crucial for accelerating drug discovery [2]. Most drug targets are cellular proteins that selectively interact with small molecule compounds for therapeutic or diagnostic purposes [18]. Existing deep learning-based DTI models usually take compound SMILES and protein amino acid (AA) sequence as input, predicts their binding label with diverse feature encoding schemes and various state-of-the-art architectures.

DeepDTA [19] utilizes drug and target sequence information for drug-target affinity prediction. It employs convolutional neural networks (CNNs) to encode features from protein sequences and compound SMILES strings and then concatenates them to predict the affinity score.

DeepConv-DTI [20] applies CNN with varying window sizes to detect local residue patterns from protein sequences, and represents drugs with extended-connectivity fingerprint (ECFP) which shows potential for binding site prediction.

DeepPurpose [21] provides several combinations of different protein encoding modules and compound encoding modules for DTI prediction. In this study, we choose message-passing neural network (MPNN) for chemical graph feature extraction and CNN to distillate protein features from AA sequence decomposition.

MDeePred [22] incorporates multiple types of protein features, including evolutionary and physicochemical properties, to enhance the drug-target affinity prediction.

GIFDTI [23] combines Convolutional Neural Networks (CNNs) and Transformers to capture long-range relationships between atoms or amino acids. It employs additional modules to independently extract global molecular features and inter-molecular interaction features for DTI prediction.

PerceiverCPI [24] utilizes sub-attention mechanisms and cross-modal attention mechanisms to identify subsequences and substructures within drug molecules and between drug molecules and protein sequences that are useful for DTI prediction.

Although existing deep learning-based DTI prediction models have demonstrated promising results, our preliminary study revealed that their prediction outcomes exhibit inconsistency. This inconsistency arises from the diversity in model architectures and the different aspects they emphasize regarding the drug-protein relationship. To address these challenges, we propose our Mixture-of-Experts (MoE) architecture, EnsDTI, which is introduced in the next section.

## 3 METHODS

The overall pipeline of our MoE model, EnsDTI, is depicted in Fig. 2. The total training and testing process could be separated into four stages: (I) train experts; (II) generate experts answers; (III) train MoE, and (IV) calculate confidence of prediction. illustrating the complete process of boosting and stabilizing current DTI prediction experts with our method. In this section, we will describe in detail: (1) how we train EnsDTI with DTI expert models (stage I to III), and (2) how we use inductive conformal prediction (ICP) to provide confidence values for prediction (stage IV).

**Figure 2:**
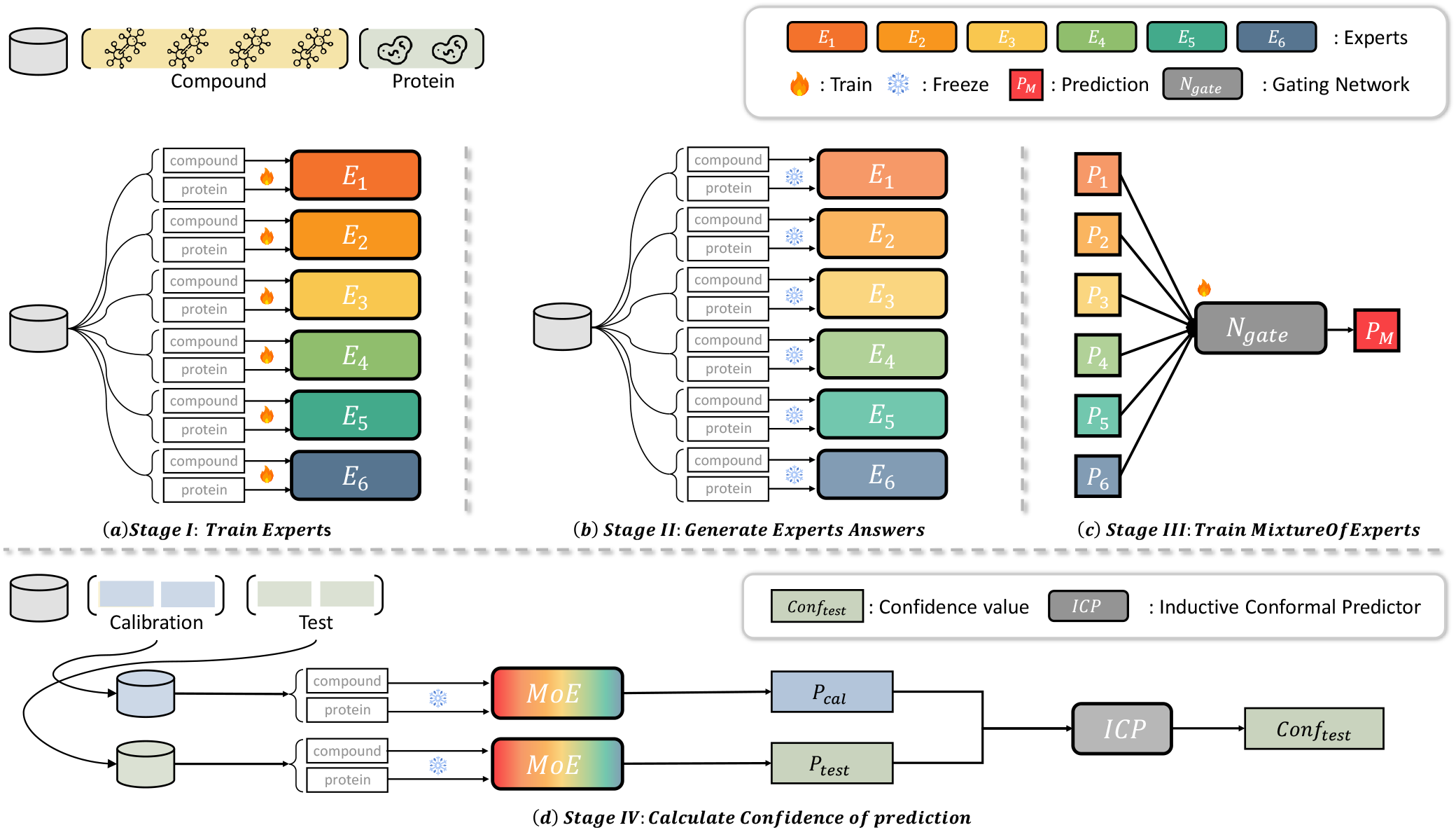
Overview of EnsDTI: (a) In the first stage, we re-train the DTI experts on each database; (b) after training, the parameters of all experts are fixed, and their predictions are generated for mixture-of-experts (MoE) training; (c) the MoE prediction is obtained by training a gating network using the experts’ predictions as input; (d) to further calibrate the predictions, we apply an inductive conformal predictor on top of the MoE to calculate the confidence value for each sample.

### 3.1 Problem Formulation

Given a pair of drugs *d* ∈ *D* and a protein *p* ∈ *P*, the objective of DTI task is to predict their binding label *y* ∈ *Y, Y* = {‘*binding*’, ‘*non-binding*’}. In the context of deep learning-based DTI models, a non-linear function *f* : *D* × *P* → *Y* is constructed to predict binding labels from *Y* of drug-protein pairs from *D* and *P*. Hence, we denote a set of DTI expert models as *F* : *f*_1_, *f*_2_, …, *f*_*n*_. In this work, we propose to mix the experts with a gating network, denoted as *g*, which operates on the existing functions *F*, to map from *F* to *Y*. This can be written as *g*(*f*_1_, *f*_2_, …, *f*_*n*_) → *Y*.

### 3.2 EnsDTI: Mixture of DTI experts

Mixture of experts (MOE) is a powerful machine learning technique that leverages multiple specialized models to enhance overall prediction accuracy and robustness. By integrating the strengths of various models, MoE effectively captures different aspects of the data, leading to improved performance. This approach also mitigates overfitting and reduces the risk associated with relying on a single model, which may have inherent limitations or biases [25, 26].

In constructing EnsDTI, we utilized six state-of-the-art (SOTA) DTI experts: DeepDTA, DeepConv-DTI, DeepPurpose, MDeePred, GIFDTI, and PerceiverCPI. As illustrated in Figure 2 (a), each DTI expert was trained with the reported optimal parameters using a 5-fold cross-validation setting. Upon completion of the training process, the parameters of all experts were fixed. We then generated soft label predictions from these models, representing their confidence in the interaction between a given compound and protein pair (Figure 2 (b)).

To further enhance the model, we introduced a multilayer perceptron (MLP)-based gating network *g* that maps from *F* to *Y* (Figure 2, (c)). Formally, this can be represented as follows:

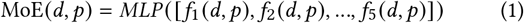

Consequently, EnsDTI incorporates the insights from individual DTI experts to enhance the overall predictive performance.

### 3.3 ICP-based Confidence Measurement

However, no model can accurately predict all samples. Providing confidence levels for each prediction based on the MoE model can further assist researchers in identifying small molecules that are likely to bind to target proteins. The ICP framework, which independently estimates the credibility of model predictions, can be readily applied to assess the confidence levels of each sample [27]. For each test sample, ICP predicts the likelihood of assignment to each available class by comparing its prediction with those from the training dataset.

Therefore, we incorporate the ICP framework [27] into EnsDTI to assess the confidence levels of model predictions. Specifically, let *X* denote the full dataset comprising the drug set *D*, target set *P*, and their interactions *Y*. ICP divides *X* into three subsets: the training set *X*_train_ used for model training, the calibration set *X*_calib_ with known labels, and the test set *X*_test_ with unknown labels. The confidence value for each test sample is then measured by comparing the non-conformity score *α*_*i*_ of the test sample to the non-conformity scores of the calibration samples in *X*_calib_.

The non-conformity score *α*_*i*_ for each sample *x*_*i*_ from either *X*_calib_ or *X*_test_ is calculated as follows:

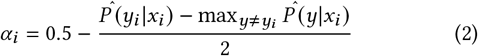

where 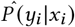 is the predicted probability of the true label *y*_*i*_ given the input *x*_*i*_. The non-conformity score measures the dissimilarity between each test prediction and the predictions on the calibration set. This calculation is performed for each possible label *y*_*i*_ in *Y* according to Equation 2, evaluating the suitability of classifying the test sample into each corresponding class.

Finally, to determine the prediction with the highest confidence, we compute the *p*-value as:

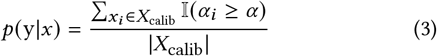

where *α* represents the non-conformity score of the test sample *x*. Each test sample is then assigned to the class with the highest *p*-value, indicating the predicted label with the greatest confidence.

## 4 EXPERIMENTS

### 4.1 Data Pre-processing

To fairly evaluate EnsDTI with baseline models, we use four benchmark databases that generated from different experiment settings to make sure each database has a distinct distribution of data.

- **davis [28]** contains data from kinase selectivity assays with their relevant inhibitors measured in *K*_*d*_.
- **kiba [29]** is constructed by combining data with different types of bioactivity (*IC*_50_, *K*_*i*_, and *K*_*d*_).
- **kinome [22]** is generated from ChEMBL (v25) by filtering representative compounds.
- **human [30]** contains interactions across different species with the same number of highly credible negative samples of each compound-protein pair.

To adapt the data for our task and ensure that the data can be utilized by each model, we removed compounds with non-covalent bond, and binarized the affinity values according to [29]. Detailed information (e.g. the positive-negative ratio, number of samples) about the data is listed in the Table 1.

**Table 1:**
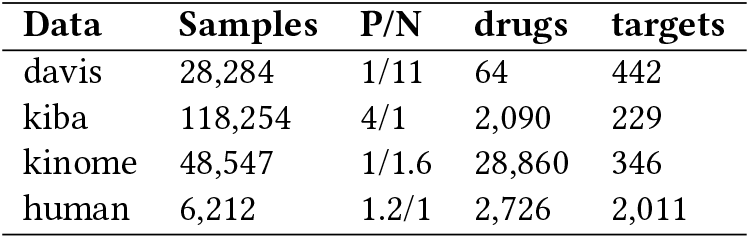
Databases used for cross-validation model evaluation. P/N: refers to the positive:negative ratio of the dataset.

| Data | Samples | P/N | drugs | targets |
| --- | --- | --- | --- | --- |
| davis | 28,284 | 1/11 | 64 | 442 |
| kiba | 118,254 | 4/1 | 2,090 | 229 |
| kinome | 48,547 | 1/1.6 | 28,860 | 346 |
| human | 6,212 | 1.2/1 | 2,726 | 2,011 |

We evaluated EnsDTI by comparing it with the DTI expert models used in our framework under the previously mentioned DTI benchmark databases: davis, kiba, kinome, and human. Under each database, we employed the k-fold cross-validation setting for all the expert models and EnsDTI. Specifically, we randomly partition each database into k subsets and conducting experiments k times. In each iteration, one subset is designated as the test set to evaluate the model’s performance, while the remaining k-1 subsets are used for model training. By repeatedly reshuffling the data and rotating the test set among the subsets, k-fold cross-validation provides a comprehensive assessment of the model’s performance on different subsets of the data.

To ensure the validity of our model, we randomly sampled and split 10% of the training data to create a validation set. This validation set is used to tune hyperparameters and make informed decisions during model training. In this study, we set k to 5, which means the dataset was divided into five equally sized subsets, and the training and evaluation process was repeated five times, each time using a different subset as the test set.

### 4.2 Evaluation Metrics

To evaluate the performance of our model, we employed three widely used metrics in drug-target interaction prediction: accuracy (ACC), area under the receiver operating characteristic curve (AUROC), and area under the precision-recall curve (AUPRC). These metrics play a crucial role in assessing the predictive capabilities of computational models in this domain.

The ACC measures the overall correctness of the predictions. It is computed as the ratio of the number of correctly predicted interactions to the total number of interactions in the dataset. Mathematically, ACC can be expressed as:

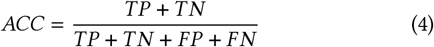

where TP represents true positives, TN represents true negatives, FP represents false positives, and FN represents false negatives.

The AUROC is a widely used metric for evaluating the model’s ability to distinguish true interactions from non-interactions across different decision thresholds. It measures an area under the ROC curve, which consists of the true positive rates (sensitivity) against the false positive rates (1-specificity) at various threshold values. AUROC ranges from 0 to 1, with higher values indicating better predictive performance. The AUROC can be calculated as:

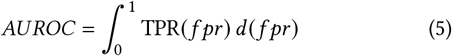

where TPR represents the true positive rate and fpr represents the false positive rate.

Another important metric for drug-target interaction prediction is the AUPRC. It provides insight into the model’s ability to prioritize true interactions over non-interactions, particularly in scenarios where the dataset is imbalanced. The AUPRC is calculated by integrating the precision-recall curve and is given by:

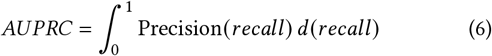

## 5 RESULTS

### 5.1 Overall Performance

#### 5.1.1 MoE architecture improves the performance of DTI prediction

We performed a comprehensive comparison between EnsDTI and existing DTI expert models under a 5-fold cross-validation setting across databases with different data distributions. As shown in Table 2, EnsDTI significantly improves the performance of current DTI models across various databases. Even when some experts report untrustworthy performance, such as in the Davis database where three out of six expert models report an AUPRC value equal to or below 0.5, EnsDTI still achieves high performance under these circumstances (AUPRC=0.670). This demonstrates its ability to leverage the strengths of each expert for more accurate predictions. The best-performing model is underlined in Table 2, highlighting that existing DTI experts excel in specific databases while performing poorly in others. Despite the varying expertise of each expert model across different databases, our MoE framework can balance their knowledge with the gating network, providing stable and robust predictions across diverse data distributions.

**Table 2:** Performance comparison of our MoE architecture (EnsDTI) and existing DTI models under cross-validation evaluation. We highlighted the models with the best performance with bold and the best performed expert model with underline text.

| Model | davis |  |  | kiba |  |  | kinome |  |  | human |  |  |
| --- | --- | --- | --- | --- | --- | --- | --- | --- | --- | --- | --- | --- |
|  | ACC | AUROC | AUPRC | ACC | AUROC | AUPRC | ACC | AUROC | AUPRC | ACC | AUROC | AUPRC |
| DeepDTA | 0.938 | 0.901 | 0.603 | 0.875 | <u>0.915</u> | <u>0.955</u> | 0.730 | 0.802 | 0.716 | 0.893 | 0.888 | 0.921 |
| DeepConv-DTI | 0.924 | 0.828 | 0.440 | <u>0.890</u> | 0.909 | 0.953 | <u>0.767</u> | <u>0.821</u> | 0.757 | 0.906 | 0.905 | 0.938 |
| DeepPurpose | <u>0.939</u> | <u>0.923</u> | <u>0.636</u> | 0.773 | 0.715 | 0.910 | 0.610 | 0.602 | 0.550 | <u>0.908</u> | 0.910 | 0.943 |
| MDeePred | 0.929 | 0.869 | 0.500 | 0.790 | 0.599 | 0.895 | 0.670 | 0.696 | 0.627 | 0.763 | 0.773 | 0.859 |
| GIFDTI | 0.927 | 0.912 | 0.604 | 0.886 | 0.914 | 0.944 | 0.742 | 0.802 | **0.857** | 0.905 | **0.970** | <u>0.976</u> |
| CPI-Perceiver | 0.928 | 0.863 | 0.488 | 0.877 | 0.901 | 0.946 | 0.714 | 0.762 | 0.826 | 0.825 | 0.904 | 0.920 |
| EnsDTI | **0.949** | **0.929** | **0.670** | **0.906** | **0.942** | **0.958** | **0.795** | **0.866** | 0.790 | **0.959** | 0.958 | **0.977** |

#### 5.1.2 EnsDTI shows stable prediction across different distribution of data when compared with voting methods

The MoE can be viewed as a specialized voting scheme that automatically learns the decision boundary for voting through the trained gating network. To evaluate whether this automated decision boundary is superior to traditional methods for aggregating predictions from DTI expert models, we compared it with conventional voting schemes, including soft-voting, hard-voting, and weighted-voting, across all databases. Specifically, the soft-voting method derives the final prediction by averaging the predicted probabilities from the six experts; the hard-voting method reports the label prediction based on the majority vote from the six experts; and the weighted-voting method calculates the final prediction by summing the weighted accuracy of each model with its prediction label.

We visualized the distribution of prediction probabilities of our MoE framework alongside these three voting methods in Fig. 3. The results indicate that traditional voting mechanisms can be significantly influenced by the results of the expert models. For instance, as shown in the top sub-figures, under both the Davis and Kinome databases, all three voting methods perform well in predicting data with positive labels but report a large number of false positives, whereas EnsDTI reports a lower number of false positives through the learned gating network. Similarly, in the left bottom sub-figure, EnsDTI reports fewer false negatives than the other methods for aggregating predictions from the DTI expert models. The only scenario where EnsDTI may underperform compared to traditional voting schemes is in databases such as Human, where most expert models already report high performance and thus make predictions with high probabilities. In such cases, our MoE framework could be influenced by some of the models, leading to interaction probabilities closer to 0.5.

**Figure 3:**
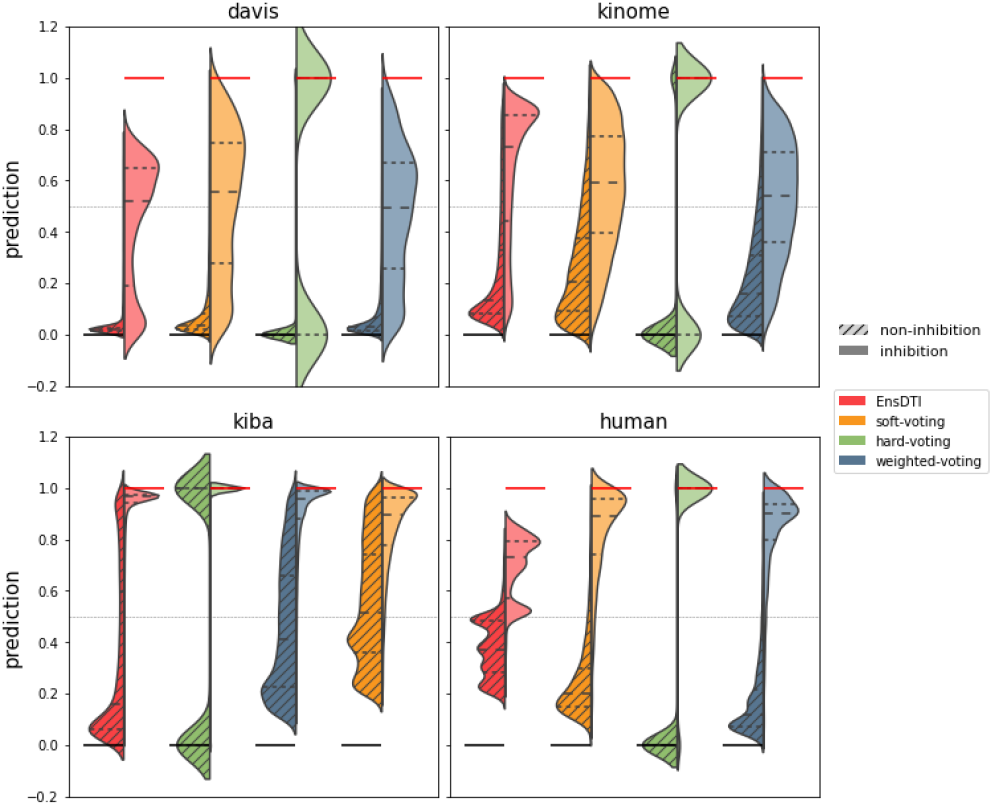
EnsDTI shows stable prediction across different distribution of data. We compare our model with voting methods to evaluate whether the MoE mechanism helps to find a better decision boundary with the knowledge of experts. Left and right portion of violin chart represent the prediction distribution of data with the true label of positive and negative respectively. Details in Section 5.1.2.

#### 5.1.3 EnsDTI outperforms DTI expert models even on the data they are confused with

For certain challenging or deceptive data points, how accurately can the mixture-of-experts mechanism ensure predictions when the majority of expert models provide incorrect answers? To intuitively assess EnsDTI’s performance on these difficult data points, we graded the difficulty of all test data and plotted the performance of each expert model and EnsDTI across different difficulty levels, as shown in Figure 4 and Figure 7. Specifically, the data difficulty level was determined based on the overall error rate of the expert models: intuitively, data points where all six experts make incorrect predictions are considered more difficult than those where only one expert predicts incorrectly. In Figure 7, the horizontal axis of each subplot represents the number of experts that correctly predicted the corresponding subset, and the vertical axis represents the accuracy performance of each model on that subset.

**Figure 4:**
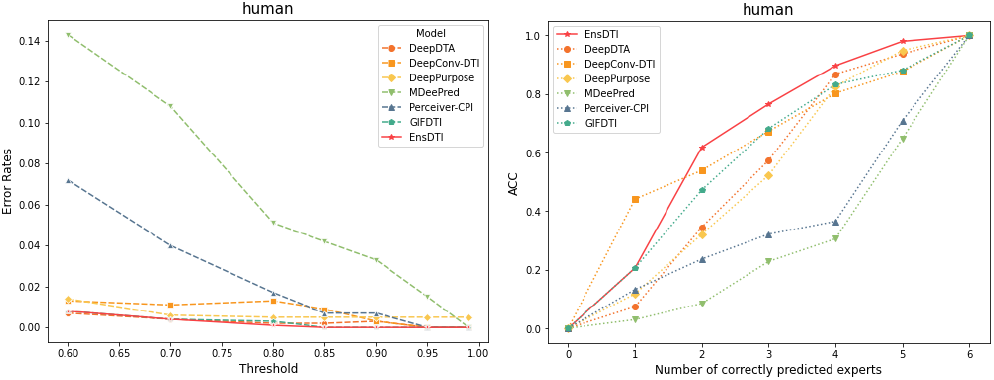
Left: Performance comparison of EnsDTI with DTI models on the difficult subsets of data. Right: Error rates under different thresholds of confidence level.

From the figure, we observe that in most cases, for the subset of samples where half of the expert models predict positive and the other half predict negative (i.e., number of correctly predicted experts = 3), EnsDTI achieves performance superior to all individual expert models. Moreover, when 4 out of 6 models correctly predict the sample, EnsDTI consistently achieves higher accuracy, surpassing 80% correctness and outperforming the other expert models. This result not only demonstrates EnsDTI’s reliability on high-difficulty datasets but also highlights the potential of incorporating more specialized experts to advance the field.

#### 5.1.4 EnsDTI Predicts DTIs with High Confidence

As described in Section ICP-based Confidence Measurement, we use the ICP mechanism to provide confidence evaluations for each prediction made by EnsDTI. We aim to determine if our model exhibits higher confidence in its predictions compared to current DTI expert models, and whether this increased confidence leads to improved accuracy. Thus, in Figure 4 and Figure 8, we plot the error rates of EnsDTI and six DTI expert models across different confidence intervals of the test subsets.

We observe that in three out of the four datasets, when EnsDTI’s confidence in its predictions is above 60%, the error rate does not exceed 1%. In the remaining dataset (kiba), EnsDTI still achieves a lower error rate than the other models, maintaining below 10%. Compared to the other models, only in the davis dataset does EnsDTI’s error rate slightly exceed that of GIFDTI and PerceiverCPI. However, for the subset of samples with confidence levels above 95%, EnsDTI still achieves the lowest error rate. Overall, as EnsDTI’s confidence in its predictions increases, the model’s error rate decreases. These results not only demonstrate that ICP effectively aids researchers in using EnsDTI for screening potential drug candidates, but also prove that EnsDTI is a more reliable choice compared to other models across most datasets.

#### 5.1.5 EnsDTI has potential in inferring target bioactivities

Although models trained on binarized labels are not designed to predict precise affinity values, their prediction probabilities can still implicitly reflect the likelihood of how effectively a small molecule can target a specific protein. To assess how well EnsDTI can infer and approximate true binding strength, we conducted the following experiment.

Using the kinome dataset, we calculated the Pearson correlation between each model’s prediction probabilities and pChEMBL values (Table 3). We utilized the standardized pChEMBL data from MDeePred [22] to indicate the bioactivity strength of compounds against their targets [31]. As shown in Table 3, existing models did not demonstrate significant capability to infer binding strength since they were trained on binarized interaction labels. However, EnsDTI’s predictions exhibited a weak positive correlation with bioactivity values, outperforming other models and indicating its potential to fit true bioactivity data and be applied in practical drug development processes.

**Table 3:**
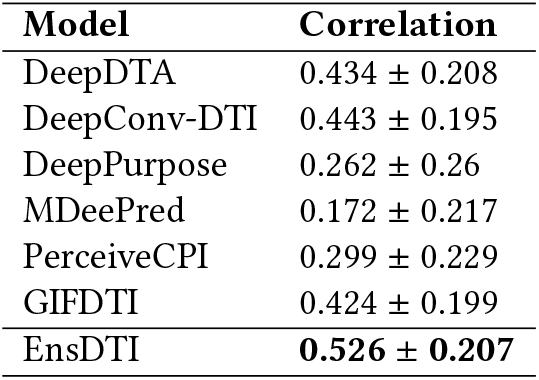
Concordance of bioactivitiy pChEMBL values with predicted probabilities of models.

**Table 4:**
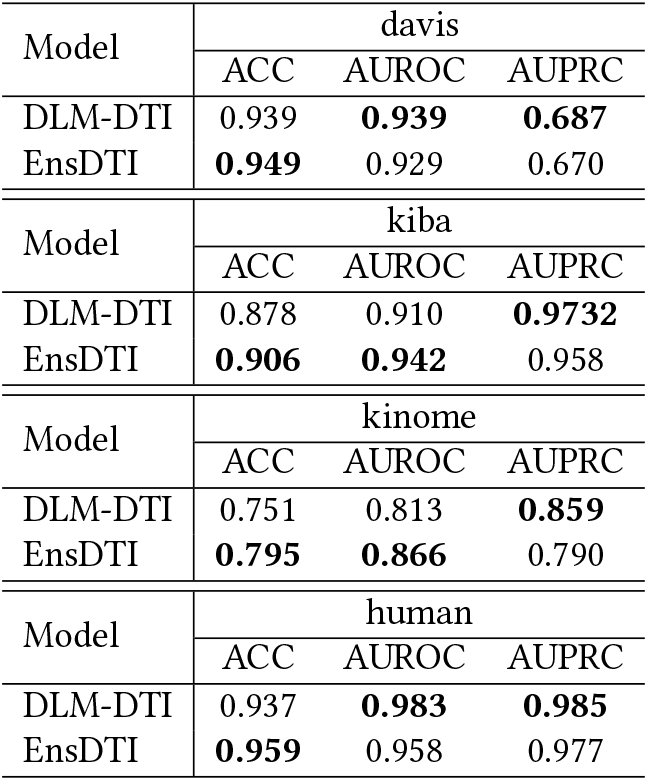
Performance comparison of EnsDTI with DLM-DTI.

We further explored whether our model can robustly fit the true binding affinities for two functionally similar proteins. Taking Ephrin type-B receptor 1 (CHEMBL5072) and Ephrin type-A receptor 4 (CHEMBL3988) as examples, both of these proteins are receptor tyrosine kinases that bind membrane-bound ephrin family ligands residing on adjacent cells. The kinome database contains interaction records of CHEMBL5072 and CHEMBL3988 with 33 and 28 small molecules, respectively, of which 22 small molecules are common to both proteins. We plotted correlation regression scatter plots between the predicted probabilities and actual affinity values for each model on CHEMBL5072 and CHEMBL3988. We compared EnsDTI with GIFDTI and DeepPurpose, the models that performed closest to EnsDTI as shown in Table 2.

From the results shown in Figure 5, it is evident that EnsDTI’s average prediction probabilities for probable candidate drugs targeting CHEMBL5072 and CHEMBLE3988 are most correlated with the experimental assay data. Other models either exhibit a weaker ability to fit bioactivity pChEMBL values or show less robustness when the target protein changes. The figure demonstrates that base classifiers tend to misclassify small molecules with high bioactivity—those more likely to be drugs—as negative samples, indicating potential losses in practical drug discovery processes. In contrast, EnsDTI, with fewer errors, showed a higher correlation with actual bioactivity data, reflecting its capability to approximate true binding affinities.

**Figure 5:**
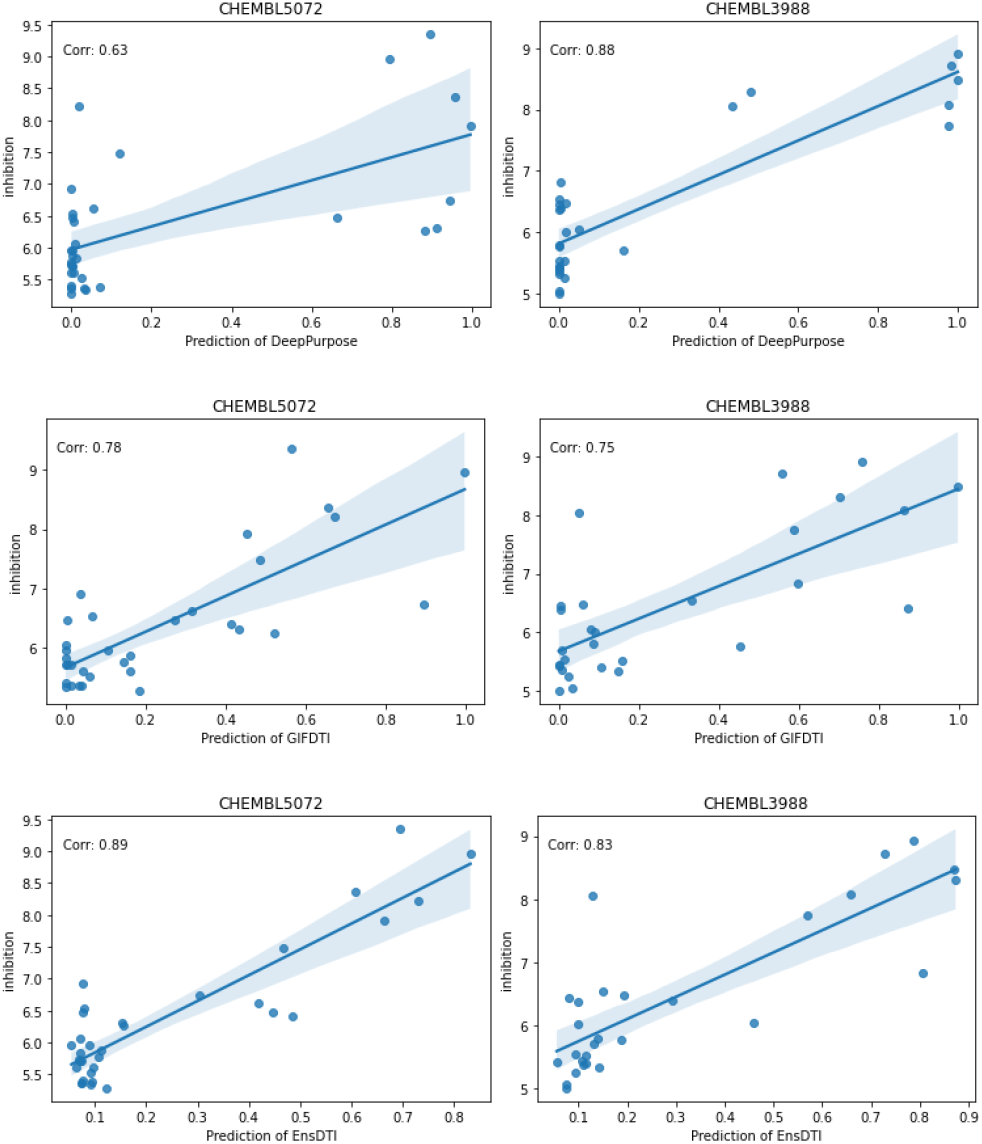
Correlation between the predicted probabilities and bioactivity pChEMBL values for each model on CHEMBL5072 and CHEMBL3988.

### 5.2 Ablation Study

In constructing EnsDTI, the number and selection of expert models are also crucial considerations. Must there be exactly six experts in the MoE framework? Which combinations can achieve comparable performance at lower training costs? These questions intrigued us. To address them, we incrementally removed each DTI expert model from EnsDTI and trained all possible ablation versions under the same experimental conditions. The complete performance table is provided in the Appendix, Table 5. Based on these experimental results, we plotted boxplots depicting the impact of the number of expert models on the performance of EnsDTI (Figure 6). The results indicate that as the number of expert models increases, EnsDTI achieves higher performance and lower bias.

**Table 5:**
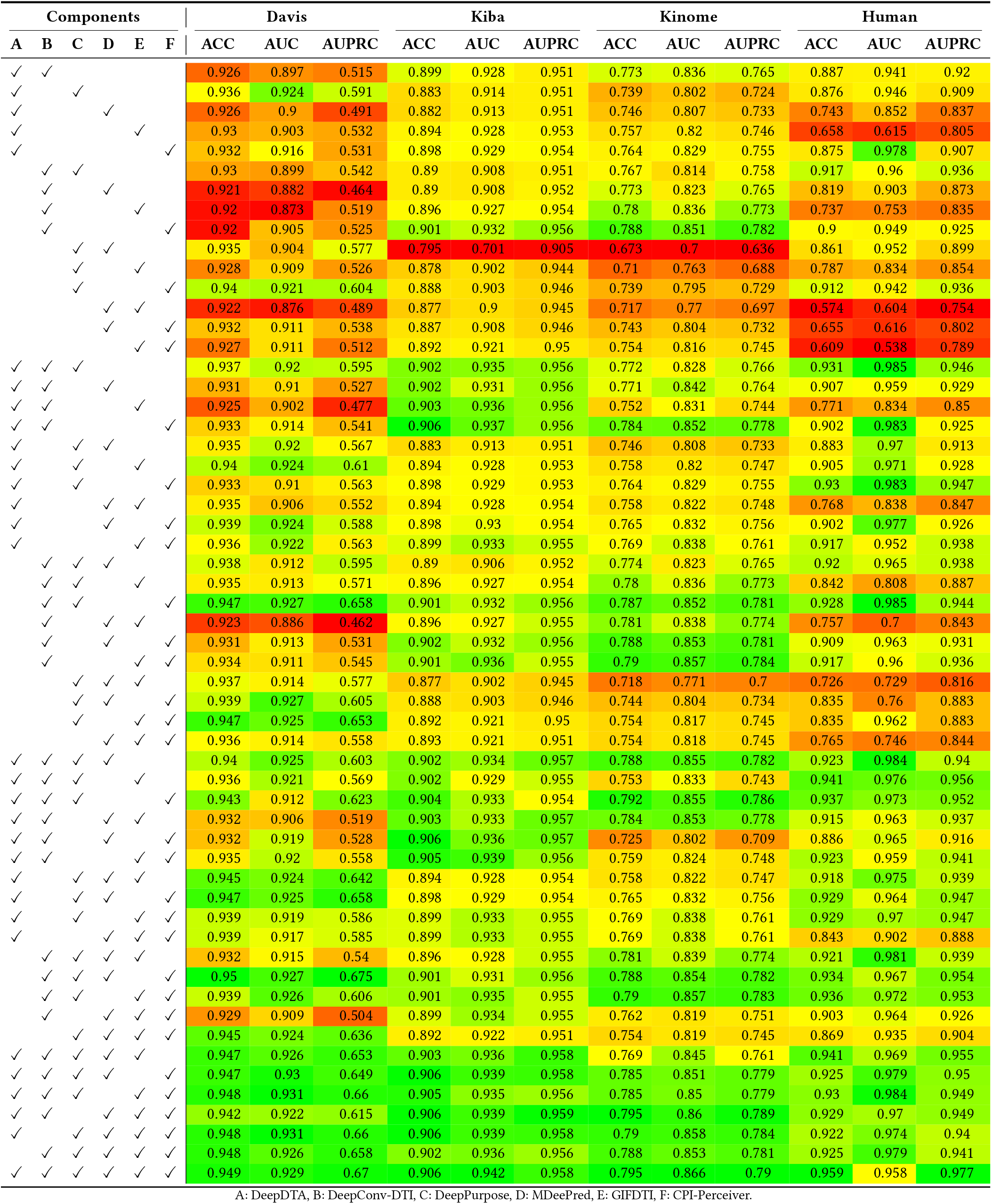
Performance of ablation study. We analyzed the performance of our model by removing the experts.

**Figure 6:**
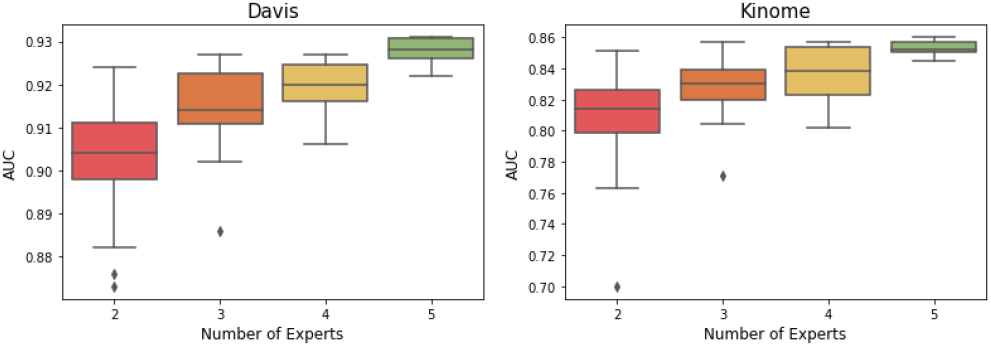
Performance of EnsDTI variants. We conducted ablation study to analysis how number of experts effect the performance of EnsDTI, details in Section 5.2.

**Figure 7:**
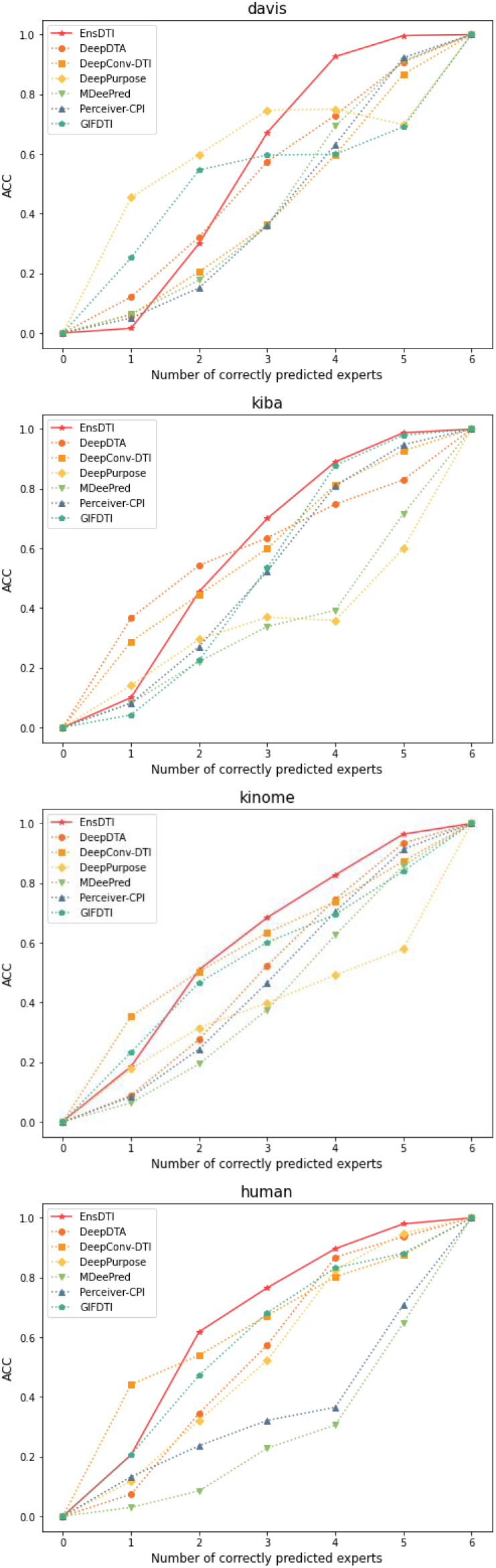
Comparison of performance on the difficult subsets of data.

**Figure 8:**
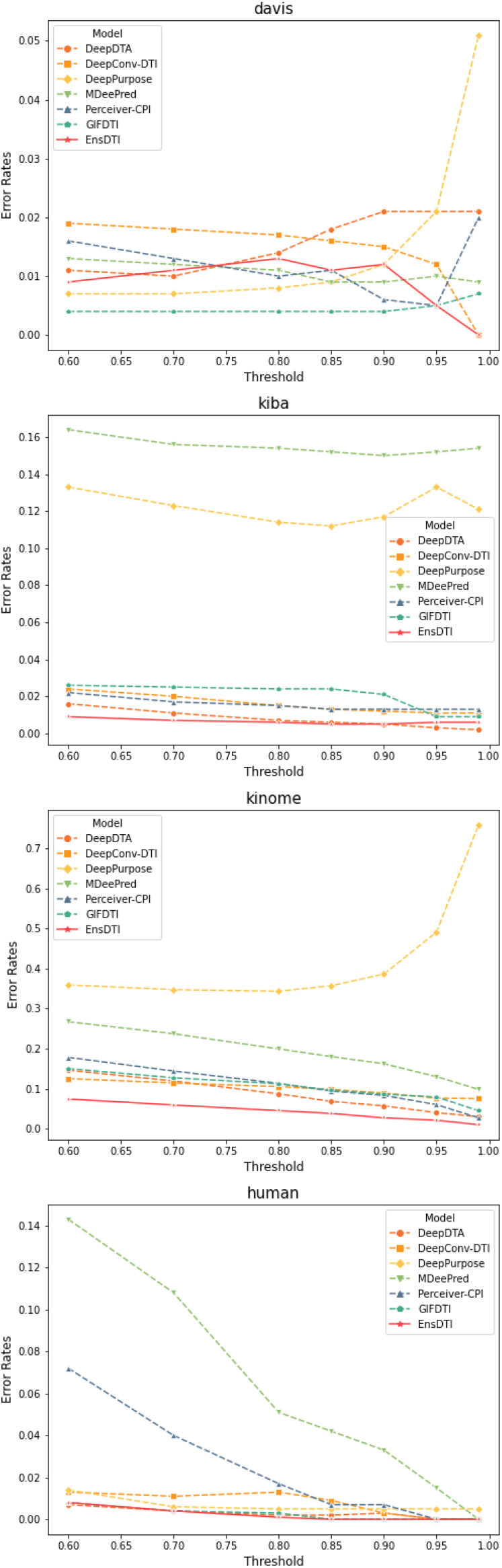
Error Rates under different thresholds of confidence level.

Additionally, the impact of the number of experts varies across different datasets: in the davis dataset, using only two experts slightly improves predictive performance but to a limited extent. Conversely, in the human dataset, using two experts to construct the MoE can negatively affect performance; for instance, EnsDTI constructed with DeepDTA and GIFDTI performs worse than each model individually. This suggests that MoE is not a universally superior method and can become confused when the individual models have significant disagreements. However, when the number of expert models in the MoE is four or more, we observed that most ablated versions of EnsDTI achieved superior predictive performance compared to individual experts.

In Table 5, cells with an orange background indicate mid-level performance among all ablation versions of EnsDTI, with colors closer to green indicating top performance and colors closer to red indicating lower performance. Furthermore, we identified some ablation versions of EnsDTI that can serve as cost-effective alternatives with minimal performance loss: in the davis and kiba datasets, EnsDTI constructed with DeepConvDTI, DeepPurpose, and PerceiverCPI; in the kiba dataset, EnsDTI constructed with DeepConvDTI and PerceiverCPI; and in the human dataset, EnsDTI constructed with DeepDTA, DeepConvDTI, and DeepPurpose all achieved commendable performance surpassing other experts.

## 6 CONCLUSION

In this work, we present EnsDTI, the first mixture-of-experts architecture for drug-target interaction (DTI) prediction, to the best of our knowledge. Our preliminary studies revealed that current DTI expert models excel in different data distributions, making it challenging for researchers to select a single model when leveraging deep learning-based approaches to narrow down drug candidates. EnsDTI addresses this issue by providing a reliable ranking list of candidate drugs for target proteins with enhanced predictive performance.

To achieve this, we propose a mixture-of-experts architecture that enhances the prediction performance of current models and incorporates an inductive conformal predictor to filter out unreliable predictions. The training process of EnsDTI can be divided into four steps: Firstly, each DTI expert model is trained to achieve its maximum performance, after which we freeze all experts to generate prediction probabilities. These prediction probabilities are then used to train a gating network, allowing the mixture-of-experts to combine the knowledge from the experts. Finally, we apply an independent inductive conformal predictor module to assign confidence values to each prediction made by EnsDTI, without interfering with the predictions themselves.

A series of empirical experiments on benchmark datasets with different data distributions demonstrated that our model effectively enhances the performance of drug-target interaction predictions compared to current DTI expert models. Moreover, the ranking list provided by EnsDTI not only has high concordance with experimental assay bioactivities but also shows consistency across different proteins, making it a reliable tool for drug-target interaction prediction.

## 7 ACKNOWLEDGMENTS

This research was supported by the Bio & Medical Technology Development Program of the National Research Foundation (NRF) funded by the Ministry of Science & ICT(NRF-2022M3E5F3085677, 2022M3E5F3085681, RS-2023-00257479), Institute of Information & communications Technology Planning & Evaluation (IITP) grant funded by the Korea government(MSIT) [RS-2021–II211343, Artificial Intelligence Graduate School Program (Seoul National University)], suppoted by Basic Science Research Program through the National Research Foundation of Korea (NRF) funded by the Ministry of Education (RS-2023-00246586) and funded by AIGENDRUG CO., LTD.. The ICT at Seoul National University provides research facilities for this study. Additionally, we extend our gratitude to Sungjoon Park, Jungseob Yi, Chang-Yun Cho, and Sangsoo Lim for their valuable discussions.

## A APPENDIX

We provide the complete results mentioned in Section 5.1.3, Section 5.1.4 and Section 5.2 in this Appendix.

Figure 7 shows the test performance (ACC) of EnsDTI and baseline models across data subsets of varying difficulty. We partitioned the original database into seven levels (indicated by the x-axis from 0 to 6, where increasing values represent higher ease and lower difficulty) based on the test results of six experts on each database. Level 0 represents the most challenging subset (where all six experts made incorrect predictions), and level 6 represents the easiest subset (where all six experts made correct predictions). We observe that, in most databases, EnsDTI has the largest ACC-#of_correctly_predicted_experts curve area and demonstrates superior performance in all subsets from levels 2 to 6.

Figure 8 shows the error rates of EnsDTI and baseline models across different confidence interval subsets. The confidence intervals are computed using ICP, and test samples with confidence values greater than 0.6 are partitioned accordingly. We observe that EnsDTI consistently exhibits lower error rates compared to the baseline models across most confidence intervals, especially when the confidence is greater than 0.9.

To further validate the effectiveness and applicability of EnsDTI, we compared it with the state-of-the-art pretrained model, DLM-DTI [32]. DLM-DTI employs pretrained ChemBERTa [33] as the drug encoder and uses pretrained ProtBERT [34] for distilling target embeddings. ChemBERTa is pretrained on 77M molecular SMILES from PubChem, while ProtBERT is pretrained on up to 393 billion amino acids. We conducted experiments on the Davis, KIBA, Kinome, and Human databases using the code and parameters provided by the authors of DLM-DTI. The dataset partitioning was consistent with that used in all other experiments. The experimental results are presented in Table 4. Our experiments demonstrate that EnsDTI achieves comparable performance to DLM-DTI, even without extensive pretraining on large datasets.

